# Neuroanatomical underpinning of diffusion kurtosis measurements in the cerebral cortex of healthy macaque brains

**DOI:** 10.1101/2020.07.25.221093

**Authors:** Tianjia Zhu, Qinmu Peng, Austin Ouyang, Hao Huang

## Abstract

**Purpose:** To investigate the neuroanatomical underpinning of healthy macaque brain cortical microstructure measured by diffusion kurtosis imaging (DKI) which characterizes non-Gaussian water diffusion.

**Methods:** High-resolution DKI was acquired from 6 postmortem macaque brains. Neurofilament density (ND) was quantified based on structure tensor from neurofilament histological images of a different macaque brain sample. After alignment of DKI-derived mean kurtosis (MK) maps to the histological images, MK and histology-based ND were measured at corresponding regions of interests characterized by distinguished cortical MK values in the prefrontal/precentral-postcentral and temporal cortices. Pearson correlation was performed to test significant correlation between these cortical MK and ND measurements.

**Results:** Heterogeneity of cortical MK across different cortical regions was revealed, with significantly and consistently higher MK measurements in the prefrontal/precentral-postcentral cortex compared to those in the temporal cortex across all 6 scanned macaque brains. Corresponding higher ND measurements in the prefrontal/precentral-postcentral cortex than in the temporal cortex were also found. The heterogeneity of cortical MK is associated with heterogeneity of histology-based ND measurements, with significant correlation between cortical MK and corresponding ND measurements (P <0.005).

**Conclusion:** These findings suggested that DKI-derived MK can potentially be an effective noninvasive biomarker quantifying underlying neuroanatomical complexity inside the cerebral cortical mantle for clinical and neuroscientific research.

## 1. Introduction

Diffusion MRI (dMRI) is a type of MRI that can quantify tissue microstructure by characterizing underlying water diffusion. With a modern scanner, a typical dMRI resolution is around 1-3 millimeters and the water diffusion detectable by dMRI is on the scale of micrometers (1). Diffusion tensor imaging (DTI) (2), a dMRI model based on Gaussian characterization of water diffusion, has been dominantly used to delineate brain tissue microstructure. Under the framework detectable with the scale of dMRI, DTI-derived fractional anisotropy (FA) (3,4) excels in quantifying well-organized microstructures in brain white matter (WM) axonal bundles and serves as a sensitive indicator of the underlying WM microstructural organization. So far most of diffusion MRI studies in the brain have been focused on white matter. DTI-derived FA can be also used to delineate microstructural architecture of the immature cerebral cortex characterized by organized radial glial scaffold in the fetal-neonate brain (5). However, cortical microstructure under more general circumstances in a brain older than fetal-neonate stage is not well organized on the scale of micrometers. FA lacks sensitivity for quantifying nearly isotropic cerebral cortex under more general circumstances and is not sensitive to cortical microstructural changes related to brain development, aging or neuropathology. Consequently, DTI-derived FA measurements cannot quantify cortical microstructure beyond the fetal-neonatal stage.

DMRI has ushered in an era in which cortical internal microstructural complexity can be studied in the brain noninvasively with more advanced dMRI modeling (e.g. 6–13) than conventional diffusion tensor model. Multi-shell dMRI is usually required for such advanced dMRI modeling. Among these advanced models, diffusion kurtosis imaging (DKI) (7) is an effective probe for quantifying cortical microstructure noninvasively. Specifically, DKI quantifies the non-Gaussian water diffusion indicative of microstructural complexity in biological tissues (14). Non-Gaussian diffusion is believed to arise from diffusion barriers such as axonal sheaths, cellular membranes, and organelles (7). Mean kurtosis (MK), a DKI-derived metric, can thus be used to quantify cortical tissue microstructure complexity and is not limited to well-organized micrometer-level diffusion environment typically found in WM.

By mapping cortical MK of preterm brains, we have revealed spatiotemporally differential cortical microstructural pattern of preterm brains and inferred regional cytoarchitectural developmental processes during that critical stage (15). Unlike DTI-derived FA that can be used to quantify cortical microstructure only in limited age range before or around birth, MK derived from DKI offers meaningful microstructural measurements in cerebral cortex of all age range in both human and animal brains (see e.g. 16,17 for review). Several studies (18, 19, 20) have shown the potential of DKI in characterizing meaningful microstructural alterations in white and gray matter across the human lifespan. Cheung et al (21) offered evidence that MK outperformed DTI-based metrics in characterizing gray matter changes for rats from birth to adulthood. DKI has been proven useful in characterizing important cerebral cortical developmental patterns as well as alterations due to brain disorders. DKI-derived MK is shown to be sensitive to cerebral cortical microstructural changes in adolescents with attention-deficit hyperactivity disorder (22). It was found that MK was significantly correlated with severity of cognitive deficiency in Alzheimer’s disease and mild cognitive impairment in both white and gray matter (23). These converging evidences suggest that DKI-derived MK is capable of quantitatively characterizing regional cortical microstructural alterations in development, aging, and disease across human and animal lifespan. Thus, regional cortical microstructure quantified by MK can be potentially used as an important biomarker for brain changes across lifespan as well as under certain diseased conditions.

Despite compelling evidence for the potential of DKI in quantitatively characterizing cortical microstructure, there have been few studies correlating DKI metrics directly with histology inside the cerebral cortical mantle. Several quantitative studies have been performed to validate DKI using histology in the WM regions (24, 25) or in brain regions including WM, cortex, and deep gray matter (26, 27, 28). However, almost no quantitative validation of DKI measurement across various regions exclusively within the cerebral cortex has been conducted so far. With each cerebral cortical region characterized by unique cytoarchitecture (29) and function, lack of precise neuroanatomical interpretation through validation makes it a challenge for cortical MK to serve as meaningful biomarkers based on noninvasive neuroimaging. Quantifying regional cortical microstructure based on histology and correlating histological measures with corresponding MK measures will fill the knowledge gap, paving the way towards “noninvasive neuropathology” by directly assessing neuroanatomical underpinning with noninvasive imaging techniques.

In this study, we aimed to test the potential roles of DKI-derived MK to be an effective noninvasive biomarker quantifying underlying neuroanatomical architecture in the cerebral cortex. We hypothesized that dMRI-based MK is heterogeneous across the cerebral cortical regions and MK is significantly correlated with neurofilament density (ND) measured based on structure tensor analysis (30) of histology images. Macaque brain model with significant cytoarchitectural differences across macaque cortical regions was used in this study. High resolution multi-shell dMRI datasets of the six post-mortem macaque brain samples were acquired. Neurofilament-stained histology images were obtained from a public website (www.brainmaps.org) (31). We explored the sensitivity of MK from dMRI to detect microstructural heterogeneity across cortical regions and demonstrated reproducibility of this microstructural heterogeneity. Linear relationship between dMRI-based MK and histology-based ND in cerebral cortex was further tested.

## 2. Methods

### 2.1 Postmortem macaque brain samples

The 6 young adult rhesus macaque (*Macaca mulatta*) brain samples were obtained by perfusion fixation with 4% paraformaldehyde after anesthesia with intramuscular injection of ketamine hydrochloride. The procedures on macaques were conducted with great care to ensure the wellbeing of the macaques and were approved by Institutional Animal Care and Use Committee at Johns Hopkins University where the procedures were performed. The macaque brain samples were obtained through cost sharing with a project for making the macaque brain atlas (32). The details about these macaques were described in our previous publication (32). The postmortem macaque brains were kept in 10% formalin for at least 2 months before high resolution multi-shell dMRI described below.

### 2.2 High resolution multi-shell dMRI and preprocessing

Before dMRI, the 6 postmortem macaque brain samples were placed in 10% phosphate-buffered saline (PBS) for at least 120 h to allow the exchange of fixation solution and PBS. The macaque brains were then transferred into a custom-made MRI-compatible container and bathed with fomblin (Fomblin Profludropolyether; Ausimont, Thorofare, NJ) for minimal background signal. Since the sizes of macaque brains are relatively big compared to rat or mouse brains, macaque brain sample and the container can be only fit into coils with very large inner diameters. Such large coils are not widely available for high magnetic field animal magnets (e.g. 9.4T, 11.7T or higher magnet field strength). Thus multi-shell (3 b values including b=0) dMRI acquisitions were conducted with a 3T Philips Achieva MR system (Philips, Best, The Netherlands) using an 8-channel knee-coil at room temperature. High resolution in this study is relevant to dMRI resolution of the 3T magnet.

Postmortem brain samples can provide high-resolution dMRI image free of the motion artifacts from in-vivo Imaging (33). However, water diffusion in the postmortem macaque brain tissue is different from that in in vivo brain for which b values for non-Gaussian diffusion are well recorded in the literature (34,35). To make sure that we captured the non-Gaussian diffusion in the cerebral cortex of the postmortem macaque brain, we acquired diffusion weighted images (DWI) at several b-values of 1500, 2500, 3500 and 4500 s mm^−2^. The diffusion signal drop of the postmortem brain gray and white matter tissue is demonstrated in Supporting Information Figure S1, which is avaliable online. Specifically, at b=4500 s mm^−2^ we could capture sufficient non-Gaussian diffusion in cerebral cortex, indicated by the deviation of the the decay curve away from a Gaussian diffusion typically observed in the ventricles. Although we could use an even higher b-value, the signal-to-noise ratio (SNR) in this high resolution dMRI limited the feasible b values to be selected. The pattern of diffusion signal drop of the postmortem brain gray and white matter tissue was reprodubicle in two test postmortem macaque brains, as also shown in Supporting Information Figure S1. Hence two b values of 1500 and 4500 s mm^−2^ were optimal for kurtosis fitting for postmortem macaque brain tissue. Two b-values (b=1500, 4500 s mm^−2^), each with 30 independent diffusion-weighted directions (36), were used to acquire coronal planes with single-shot echo-planar imaging (EPI) sequence and SENSE parallel imaging scheme (SENSititivty Encoding, reduction factor =2). High resolution dMRI parameters were: FOV=100×100×72mm^3^, in-plane imaging matrix=166×166, in-plane nominal resolution before zero-filling interpolation = 0.6×0.6 mm^2^, slice thickness=2mm, TR/TE=2100/77.8ms, and number of scan average (NSA) =24. Six b0 volumes (three b0 volumes for b 1500s mm^−2^ and thee b0 volumes for b 4500s mm^−2^) were acquired to ensure sufficient number of b0 acquisitions (36). Two repetitions were performed for each b-value acquisition to increase the SNR (37), resulting in a total acquisition time of 17 hours for scanning each postmortem macaque brain. Accordingly, datasets of 4 sessions with 2 sessions per b value 1500 or 4500 s mm^−2^ were obtained for each macaque brain.

An intra-session 12-parameter affine registration aligning all diffusion-weighted images (DWIs) to b0 image of the same session was conducted. This process was followed by intersession 12-parameter affine registrations aligning b0 image of the source session to that of the target session and the transformation matrix was applied to all DWIs in the source session. The cost-function used in the registration was mean squared intensity difference. The acquisition and preprocessing procedures above were repeated for 6 postmortem macaque brains. Unlike misalignments in *in vivo* scans with high b values (38), the misalignments across diffusion MR images caused by motion and distortion were relatively small in the imaging setting of this study where skull-stripped *ex vivo* macaque brain tissues immersed in fomblin were scanned. To demonstrate good alignment, we have included 6 Supporting Information Videos S1-S6 showing aligned diffusion-weighted images across different b-values and diffusion gradient directions for all 6 scanned macaque brains.

### 2.3 Diffusion kurtosis and tensor fitting with high-resolution preprocessed DWIs

With preprocessed multi-shell DWIs, apparent kurtosis *K_app_* and apparent diffusivity *D*_*app*_ of each diffusion gradient orientation were obtained with the following equation (7, 34):

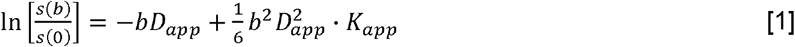

where *s*(0) is the signal without applying any diffusion gradient (i.e. b0 image), *s*(*b*) is the diffusion weighted signal intensity at two b-values (b=1500, 4500 s mm^−2^), and b is the b value. The voxel-wise MK value was calculated as the average of *K_app_* over all directions. Besides MK, axial (AK) and radial kurtosis (RK) were also calculated (35). The voxel-wise diffusion kurtosis was fitted with in-house Matlab code using constrained linear fitting. This fitting code includes Gaussian smoothing of DWIs with full width half maximum (FWHM) value of 1.5×1.5×1.5mm to remove noise (39) and improve the reliability of the kurtosis estimation (40). Although there is no “rule of thumb” for choosing FWHM for DWI image analysis, a 1.5mm isotropic FWHM size is a good balance between estimating MK at the length scale within cortical mantle and reducing noise, given macaque cortical mantle thickness 0.5 to 3.5mm (41). The procedures of DKI data acquisition and MK computation are illustrated in the left panel of Figure 1.

**Figure 1:**
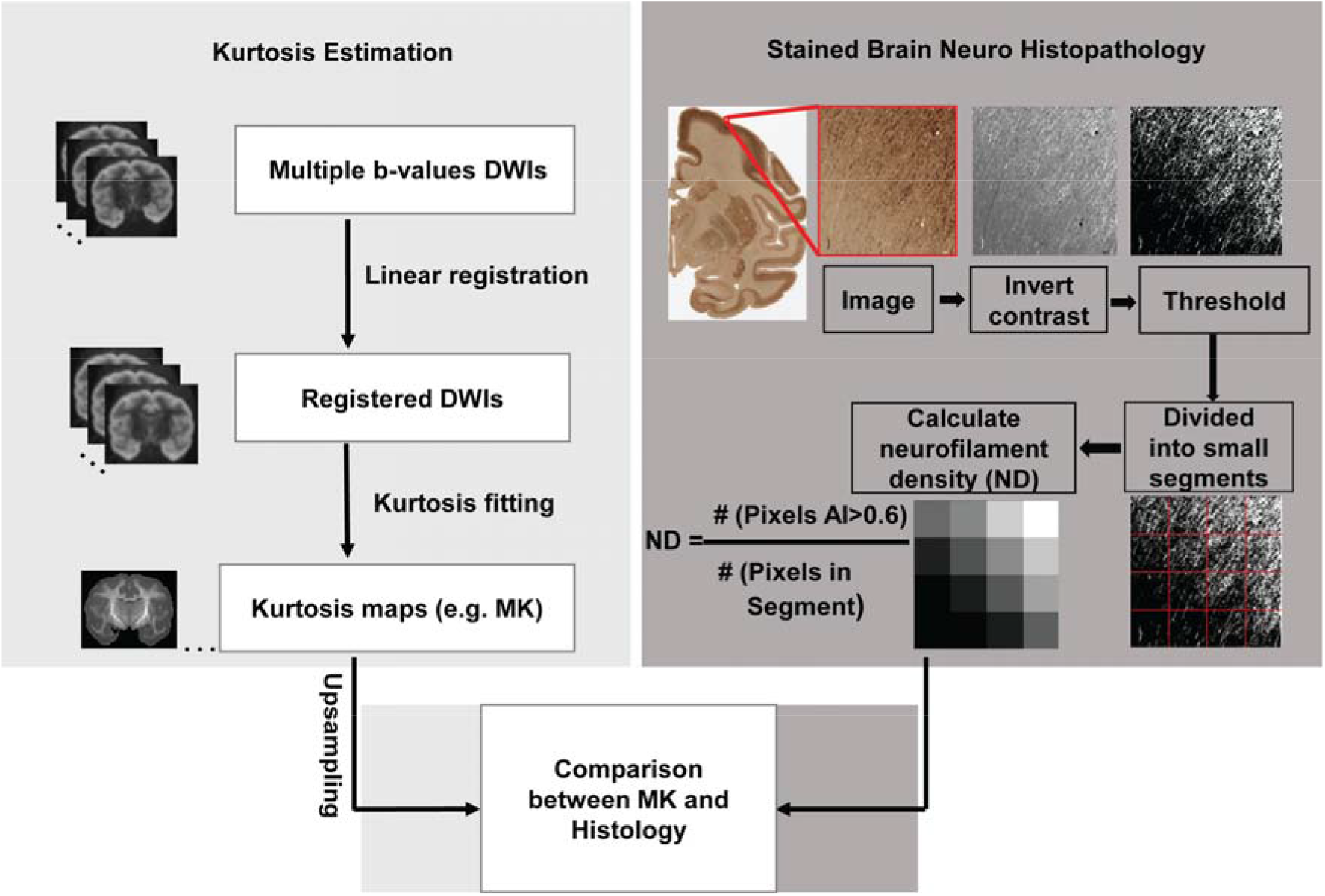
Flowchart of evaluating relationship between mean kurtosis (MK) and histology in the cerebral cortex. The flow chart consists of three technical components, estimation of MK maps in the left panel, calculation of neurofilament density (ND) in the right panel and systematic and quantitative comparison between MK and ND map. For kurtosis estimation in the left panel, the procedures included from top to bottom acquisition of diffusion weighted images (DWIs) with multiple b-value, linear registration of DWIs to the b0 image, constrained linear kurtosis fitting with registered DWI volumes to generate kurtosis maps including MK maps and upsampling of MK maps. For ND estimation in the right panel, the procedures included contrast inversion, calculation of anisotropy index (AI) for each pixel with structure tensor analysis, thresholding, blocking histological image at the similar resolution to upsampled diffusion MRI, and calculation of ND with the equation.

Diffusion tensor fitting was conducted with DWI of b 1500 s mm^−2^ besides b0 by using DTIstudio (42). Parametric maps included FA, mean diffusivity (MD), axial diffusivity (AD) and radial diffusivity (RD) derived from diffusion tensor. Averaged DWI (aDWI) maps were generated by averaging DWIs of b 1500 s mm^−2^. In this study, only MK and FA measurements at different cortical regions were assessed.

### 2.4 Neurofilament histological images of the macaque brain

High-quality macaque brain histological images of around 55,000dpi (31) were downloaded from a public website (www.brainmaps.org) to examine the neuroanatomical underpinning of diffusion kurtosis. Multiple histological images with different stains (e.g. Nissl, SMI-32 and Weil) are available from this website. Diffusional MK quantifies complexity of diffusion barrier in the brain tissue (7). To match the cerebral cortical MK measurements, histological images with SMI-32 neurofilament stain were selected. Neurofilaments provide much of the cytoarchitectural support for neurons and are one of the major underlying neuroanatomical architectures contributing to diffusion MK in the cerebral cortex. The neurofilament histological images adopted in this study were stained by SMI-32, a neurofilament marker staining neuronal cell bodies, dendrites and some thick axons in the central and peripheral nervous systems. As described in the literature (31), the Glass-mounted sections of the coronal histological slices of macaque brain stained with SMI-32 were scanned at 0.46 μm/pixel by an Aperio ScanScope T3 scanner.

### 2.5 Estimation of neurofilament density from histological images

For a stained histological image showing organized architecture, structure tensor (30) analysis can be used for estimating local orientation, anisotropy, and intensity of organized architecture in the histological image. Thus, structure tensors were adopted to estimate ND with SMI-32 neurofilament histological images. A structure tensor, with details described in the literature (30), is a matrix representation of the image partial derivatives defined as the second order symmetric positive matrix:

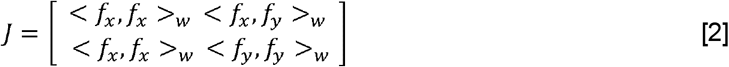

where *f_x_* and *f_y_* are the images of the partial spatial derivatives along x and y, respectively; each entry (e.g. *<f_x_, f_x_ >_w_*) is the inner product weighted by a Gaussian weighting function *w*. Structure tensors can characterize anisotropic cerebral cortical fiber structures in the scale of sub-micrometers in the neurofilament histology images with fiber orientation and anisotropy index (AI) (30). The AI, similar to DTI FA parameter, was computed by:

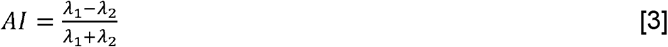

where *λ*_l_ and *λ*_2_ are the largest and smallest eigenvalues of *j*. With AI measurements from the structure tensor, fiber-like neurite microstructure in the cerebral cortex can be segmented from the background for ND measurement.

Detailed procedures of histology image analysis are illustrated in the right panel of Figure 1. For measuring ND based on histological image, a structure tensor was computed for every pixel in the histological image. Pixels with AI>0.6 were classified as fiber structure. To preserve histological details, the histological images of 55,000dpi (resolution around 0.46µm) were blocked into segments of 512×512 pixels with each segment size of 0.24×0.24mm. The DKI and DTI images (0.6×0.6 mm^2^ before zero filling) were upsampled to have exactly same 0.24×0.24mm in-plane resolution as that of blocked histological segments. In each blocked segment, the ratio of the area classified as fiber structure to the blocked area was used to estimate ND of stained histolsogical image. That is, ND was defined as the number of pixels with AI>0.6 divided by the total number of pixels in the segment. In this way, ND map with identical resolution to that of MK map was obtained for group comparison and correlation between MK and ND.

### 2.6 MK and FA differences between frontal and temporal cortices

Consistent regions-of-interests (ROIs) at frontal and temporal cortices were selected in MK or FA maps of all 6 macaque brain samples as well as ND maps from the histological data. Student t-tests were conducted to compare MK, FA or ND measurements between frontal and temporal ROIs. Bonferroni correction was conducted for multiple comparisons of MK or FA measurements across the 6 samples. Each histological slice has thickness of 0.03mm and the slice gap of histological images is 0.67mm. Thus, one MRI image with 2mm slice thickness corresponds to three consecutive histological slices. To match thickness between MRI and histological images (43), corresponding ROIs in one MK map and three consecutive ND maps were used for MK and ND measurements.

### 2.7 Correlation between ND and MK

To delineate quantitative relationship between ND and MK, MK map was reoriented to be aligned with the histological image. Corresponding slice of reoriented MK map and neurofilament histological image was chosen based on visual inspection of matched anatomical features. As elaborated in 2.6, to match thickness between MRI and histological images (43), we measured MK in one MK map and ND in three ND maps for correlation between MK and ND. With distinguished higher and lower MK values observed in the frontal and temporal cortices, 8 ROIs covering both frontal and temporal cortical areas were selected at corresponding locations in the MK map and ND maps, respectively. MK at each ROI was measured as the mean of MK of all pixels in this ROI in the MK map. ND at the corresponding ROI in three consecutive ND maps was measured as the mean of ND of all pixels in this ROI in the three ND maps. Pearson correlation was performed to test if there was a significant correlation between the MK and ND measurements at corresponding ROIs.

## 3. Results

### 3.1 High-resolution maps of DTI- and DKI-derived metrics

With high-resolution dMRI, high-resolution maps from DTI-derived metrics and DKI-derived metrics from a representative macaque sample in a coronal view were displayed in Figure 2. These maps included FA, color-encoded FA, aDWI, AD, RD, MD, MK, AK, and RK. Specifically, DKI-derived metric maps demonstrate high resolution and high SNR to delineate microstructure of the cerebral cortex, despite relatively smaller size of macaque brains compared to that of human brains. Figure 3 showed the coronal slices of MK maps from anterior to posterior brain. High contrasts between white and gray matter are clear in these MK maps. The MK value in most of white matter is above 2 while MK value in the cortex is much lower, typically from 0.5 to 1.8. Within the cerebral cortex, prominent MK contrast was observed in the coronal slices between the temporal cortex and the prefrontal/precentral-postcentral cortex, indicated by white arrows. A generally higher MK values in the prefrontal/precentral-postcentral cortex and lower MK values in the temporal cortex across these coronal slices are apparent. As appreciated from the intensity-encoded color, MK values at prefrontal/precentral-postcentral cortex can be about 2-3 times larger than those at the temporal cortex, suggesting microstructural differences between functionally distinctive cortical areas.

**Figure 2:**
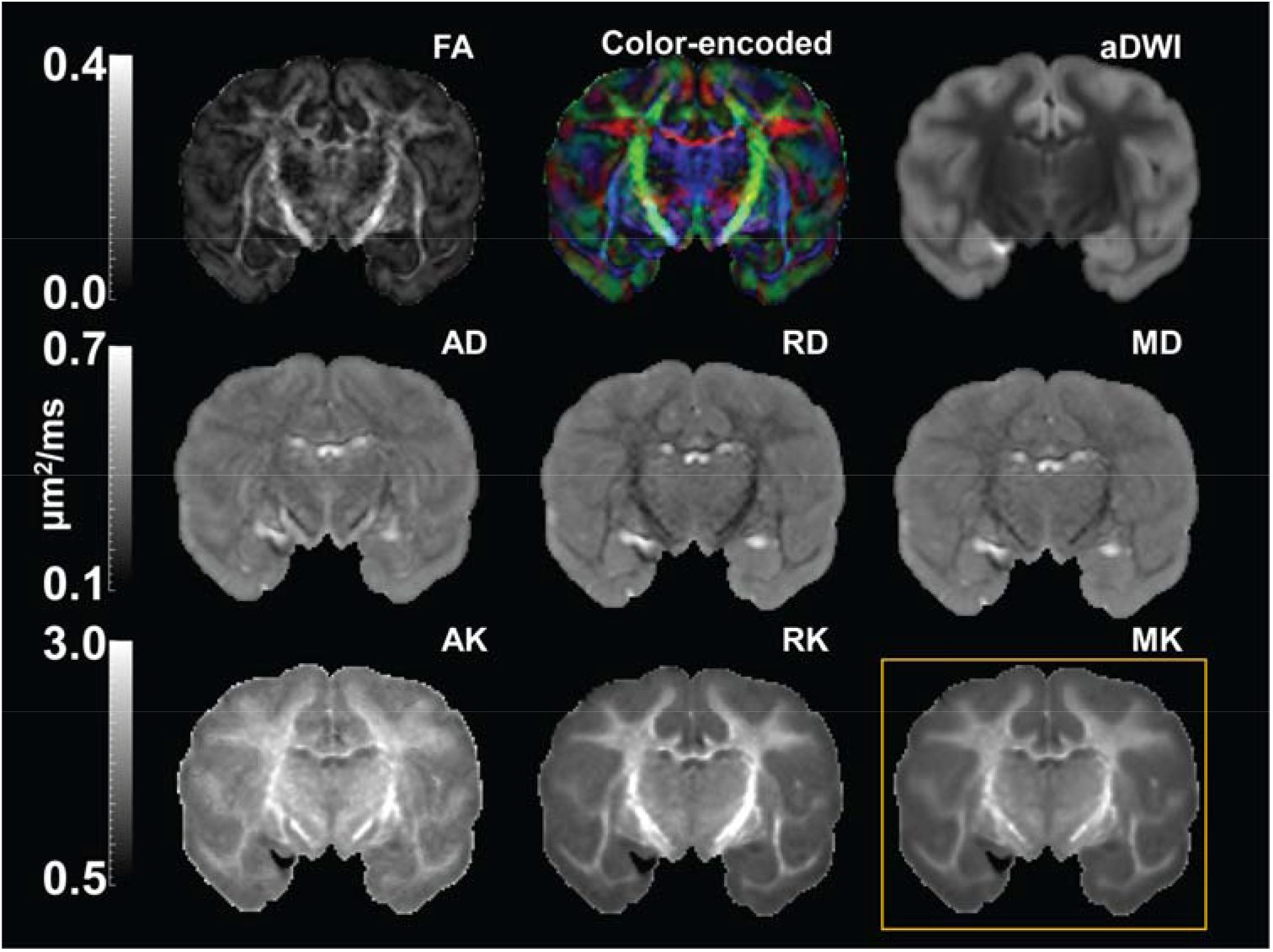
High resolution diffusion tensor imaging (DTI) and diffusion kurtosis imaging (DKI) maps from a representative macaque brain (sample #2). First two rows show DTI parameter maps, including fractional anisotropy (FA), color-encoded, mean diffusivity (MD), axial diffusivity (AD) and radial diffusivity (RD) map, along with the averaged diffusion-weighted image (aDWI). Third row shows DKI parameter maps, including axial kurtosis (AK), radial kurtosis (RK), and mean kurtosis (MK) maps. MK map shown in the yellow box was used for comparison with histology.

**Figure 3:**
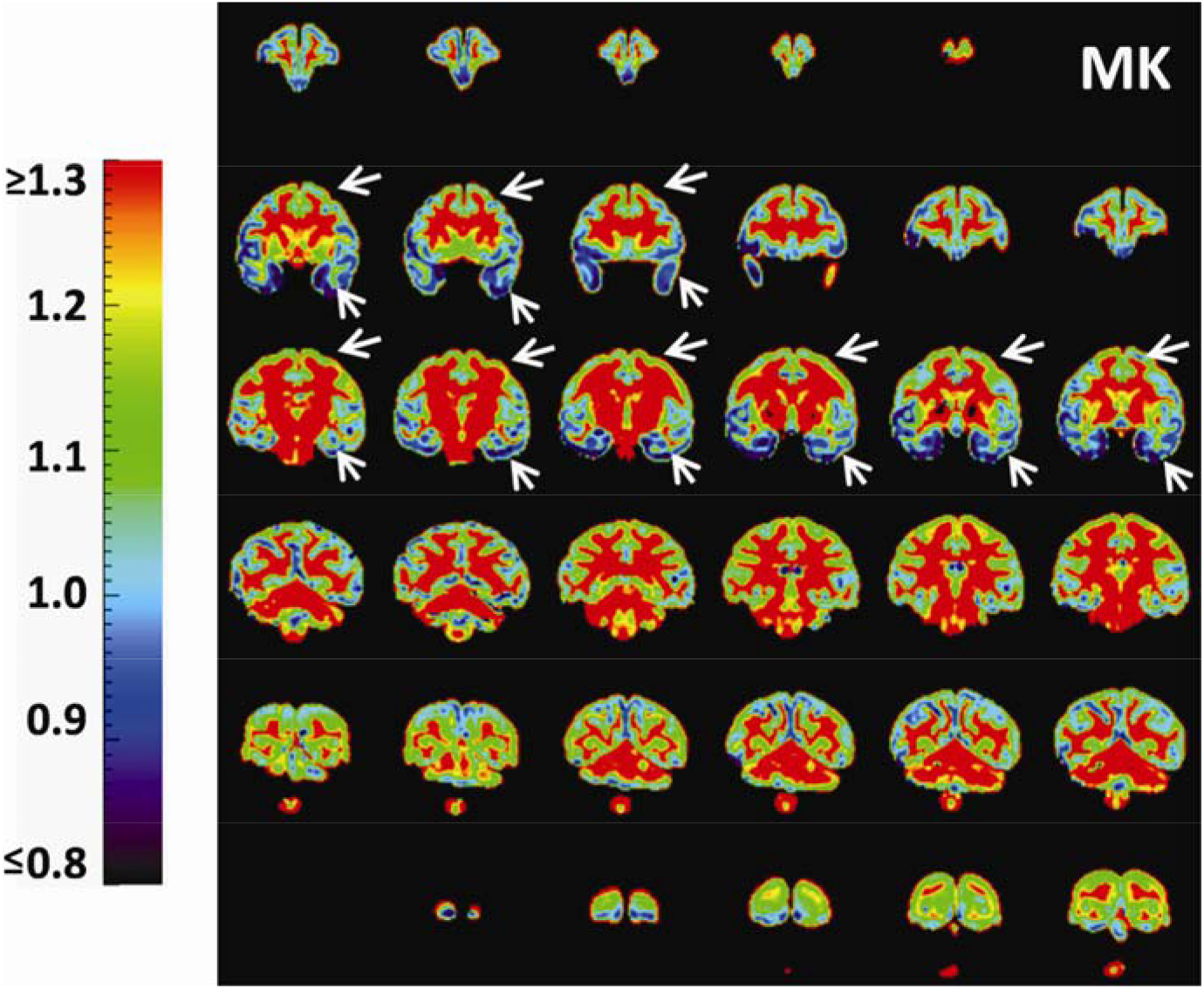
Coronal MK maps from anterior to posterior from a representative macaque brain (sample #1). Significant MK difference is clear between the temporal cortex (blue regions pointed by white arrows) and prefrontal/precentral-postcentral cortex (yellow-green regions pointed by white arrows). The color bar encodes MK values.

### 3.2 Reproducibly heterogeneous MK measurement from dMRI and ND measurements from histology across cortical regions

As shown in Figure 4, relatively higher MK at the prefrontal/precentral-postcentral cortex (indicated by the red arrow) and relatively lower MK at temporal cortex (indicated by the yellow arrow) observed in Figure 3 was reproducible across all 6 scanned macaque brains. Such heterogeneous cortical MK pattern is also consistent to neurofilament histological image. In Figure 5, denser staining in neurofilament histological images and corresponding brighter ND in ND maps at the prefrontal/precentral-postcentral cortex than at the temporal cortex are apparent across several coronal planes. Figure 5 shows that the prefrontal/precentral-postcentral cortex (indicated by purple arrows) is characterized by relatively higher ND whereas the temporal cortex (indicated by yellow arrows) is characterized by relatively lower ND. MK and ND measured at corresponding ROIs at the prefrontal/precentral-postcentral and temporal cortex in a corresponding coronal plane across all 6 scanned macaque brains are shown in Figure 6a. Significant differences (P<0.00005, Bonferroni corrected) of MK measurements between the prefrontal/precentral-postcentral cortical ROI (blue) and temporal cortical ROI (yellow) are apparent across all 6 macaque brains, further confirming reproducible heterogeneity pattern of MK measurements from dMRI. Accordingly, significantly higher ND (P<0.00001) values were found at the prefrontal/precentral-postcentral cortical ROI (blue) than at the temporal cortical ROI (yellow), consistent with the MK findings from dMRI. In comparison, there are no consistent differences between FA values in the same prefrontal/precentral-postcentral and temporal cortical ROIs across the six macaque brains, as shown in Figure 6b. Specifically, there are no significant FA differences in sample #3, #4 and #5. The significant FA differences in samples #2, #6 and sample #1 point to different directions. FA values are generally low due to lack of sensitivity of FA in nearly isotropic cortical regions.

**Figure 4:**
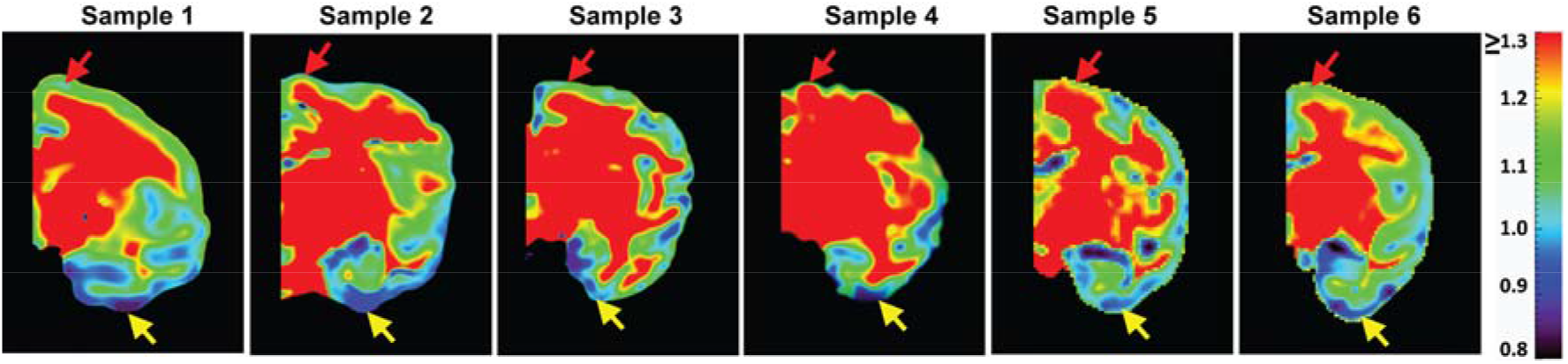
A representative MK map in a corresponding coronal slice for each of all six postmortem macaque brains included in this study. Relatively higher MK at the prefrontal/precentral-postcentral cortex (indicated by red arrow) and relatively lower MK at the temporal cortex (indicated by yellow arrow) was reproducible across all 6 scanned macaque brains.

**Figure 5:**
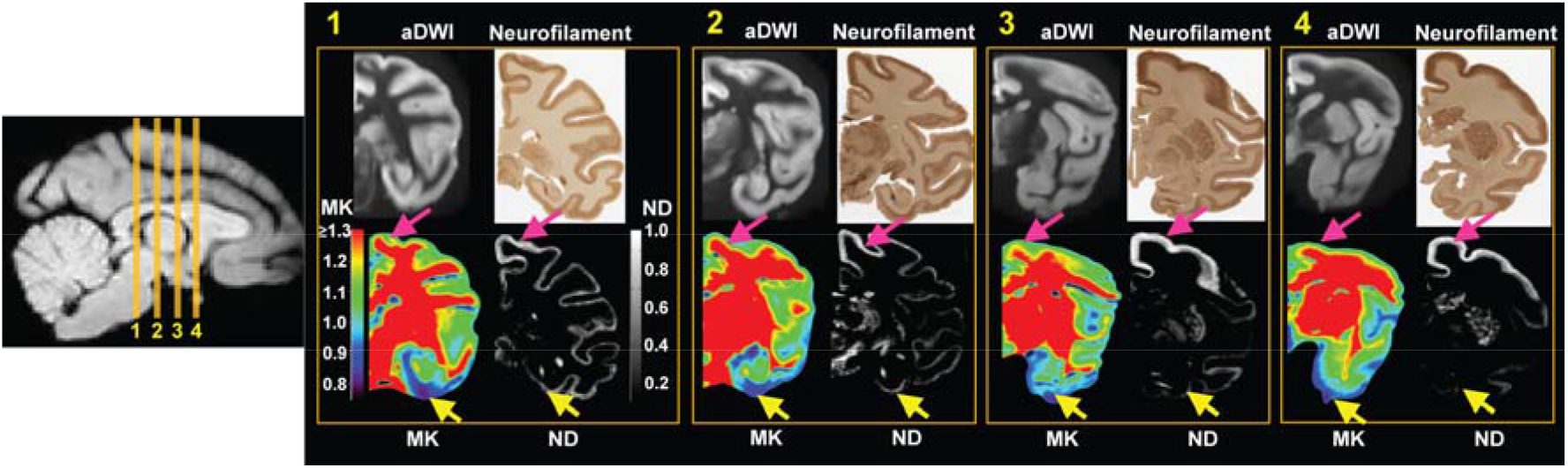
Qualitative comparison between MK and ND map in four different coronal slices 1, 2, 3 and 4 from a representative macaque brain (sample #2). In each panel, top left, top right, bottom left and bottom right shows aDWI, corresponding histology image, MK map and ND map, respectively. aDWI is shown as an anatomical reference. High MK value in the prefrontal/precentral-postcentral cortical regions is consistent to high ND value at corresponding locations indicated by pink arrow, while low MK value in the temporal cortical regions is consistent to low ND value at corresponding locations indicated by yellow arrows. The color bar and grayscale bar encode MK and ND values, respectively.

**Figure 6:**
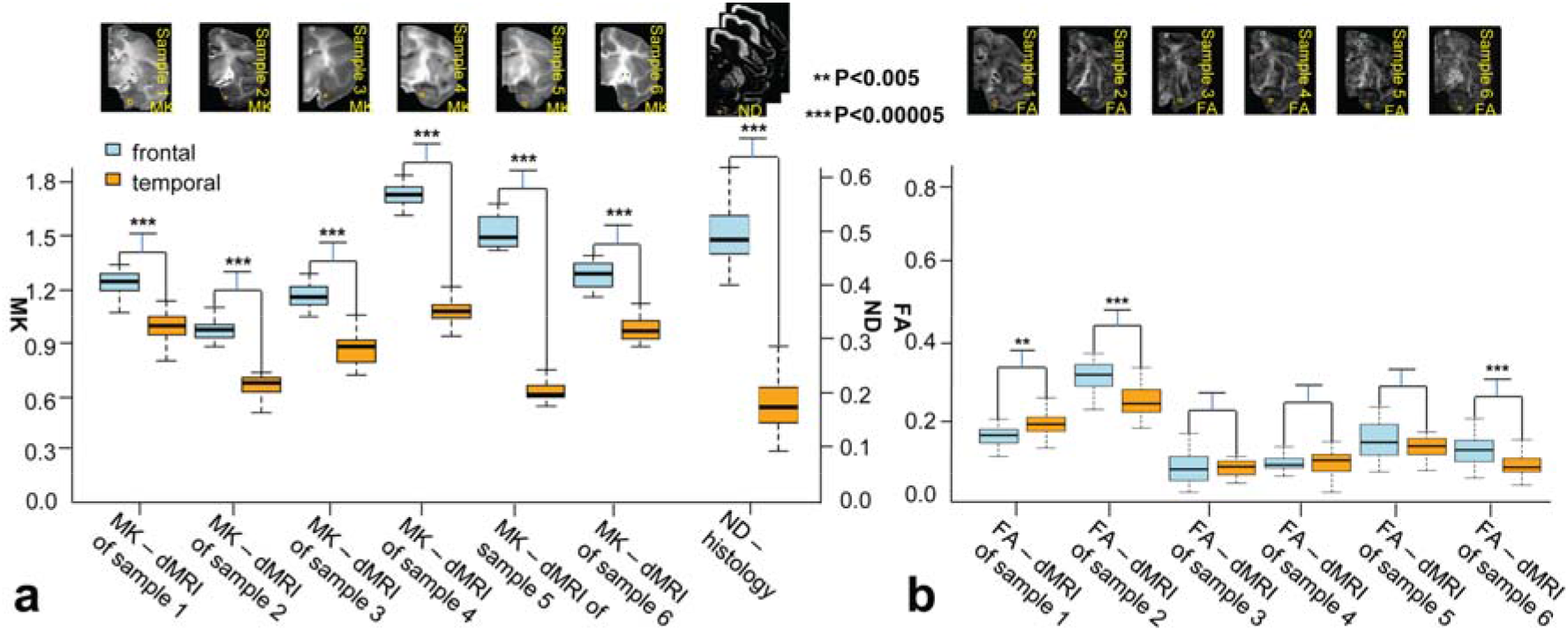
a. Reproducibly significantly higher (P<0.00005, Bonferroni corrected) cortical MK measurements at regions of interest (ROI) in the prefrontal/precentral-postcentral cortex than cortical MK measurements at ROI in the temporal cortex across all 6 macaque brain samples. MK measurements from ROI in the prefrontal/precentral-postcentral and temporal cortex were plotted for each sample as blue and orange boxplots, respectively. To match the thickness of MK coronal plane, ND measurements from the corresponding ROI on three histology slices were plotted alongside. Significant higher (P<0.00001) ND measurements in corresponding prefrontal/precentral-postcentral cortex than ND measurements in corresponding temporal cortex, shown on the right, were consistent with MK measurements. b. Cortical FA measurements at the same ROIs in the prefrontal/precentral-postcentral and temporal cortex across all 6 macaque brain samples showed no consistent differences between FA values in different cortical ROIs.

### 3.3 Significant correlation between MK and ND

Significant correlation between MK measurements at a representative coronal slice and ND measurements in three corresponding consecutive histological slices is shown in Figure 7. Significant MK differences between red and green boxes in Figure 7a is consistent to apparent ND differences in corresponding boxes in Figure 7b and 7e. With enlargement of the red and green boxes in the histological image in Figure 7b, denser staining in the red box (Figure 7c) compared to that in the green box (Figure 7d) can be well appreciated. Collectively, it can be clearly observed from Figures 7a-7d that denser staining in neurofilament histology image is associated with higher MK. With MK and ND measurements at the 8 corresponding ROIs demonstrated in Figures 7a (black circles), 7b (black circles) and 7e (white circles), statistically significant (P<0.005) linear correlation was found between MK and ND.

**Figure 7:**
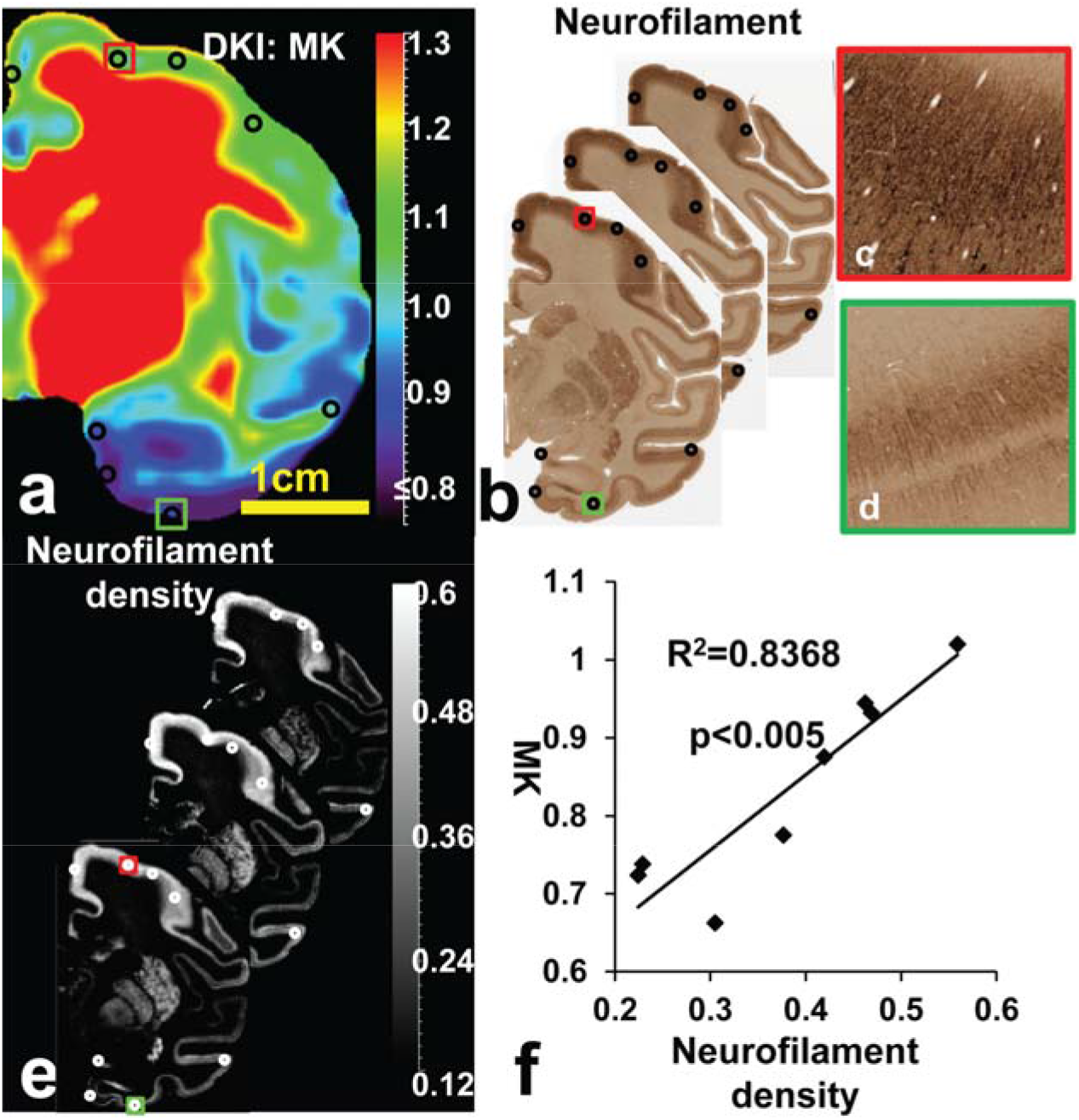
Significant linear correlation between MK and ND measurements at ROIs of a representative macaque brain (sample #1). a. MK map of a representative coronal slice. b. Corresponding three neurofilament-stained histological slices matching thickness of the MK map. c-d. Enlarged red and green ROIs show denser staining in the red box than in the green box. e. Three ND maps calculated from the three histological slices in panel b. f. With MK and ND measurements at the 8 corresponding ROIs demonstrated in panels a (black circles), b (black circles) and e (white circles), statistically significant (P<0.005) linear correlation was found between MK and ND.

## 4. Discussion

In this study, by using high resolution multi-shell dMRI and neurofilament histological images of the macaque brains, we explored the sensitivity of MK from dMRI to detect microstructural inhomogeneity across cortical regions. We further tested linear relationship between dMRI-based MK and histology-based ND in cerebral cortex. It was found that the cortical microstructural differences characterized by significant MK measure differences between prefrontal/precentral-postcentral and temporal cortex are consistent and reproducible across different brain coronal planes and across 6 macaque brain samples. So far cortical MK has only been qualitatively described as an indicator of microstructural complexity. Sensitivity of cortical MK to distinctive cortical microstructure across brain regions revealed in the presented study makes cortical MK a potentially valuable biomarker. With ND in neurofilament-stained histological images quantified by the structure tensor, significant microstructural differences across functionally distinctive cortical regions were confirmed by ND measurement. Furthermore, it was found that dMRI-based MK measurements were significantly and positively correlated to corresponding histology-based ND measurements at the ROIs in the prefrontal/precentral-postcentral and temporal cortex. We directly identified the neuroanatomical underpinning of cortical MK with histology-based measures, despite that other neuroanatomical parameters such as neuronal density may also contribute to the MK values. This study presented quantitative linear correlation between dMRI-based MK and histology-based ND across regions exclusively inside the cerebral cortex, paving the way for assessment of cortical histological measures (e.g. ND in this study) with noninvasive imaging techniques. Collectively, these findings suggested that dMRI-based MK can be an effective noninvasive biomarker quantifying underlying regional neuroanatomical architecture in the cerebral cortex.

### 4.1 Cortical microstructure characterized by dMRI

As an important *macro*structural biomarker, cortical thickness or volume across cortical regions has been extensively studied in the last two decades. However, cortical *micro*structure consisting of distinctive rich neuronal information that cannot be accessed by cortical thickness or volume measurement has been less studied. DMRI so far has been primarily used to characterize WM microstructure in the past few decades. Relatively few dMRI studies have been conducted for investigating cortical microstructure. With well-organized microstructure (e.g. radial glial scaffold) in the immature cerebral cortex of developing human (e.g. 5,15,44–51) and animal brain (e.g. 52–58), DTI-based metric measurements such as FA values can be used to infer the brain developmental pattern characterized by dendritic arborization, synaptic formation, and neuronal differentiation during the fetal-neonatal stage. After that stage, cerebral cortical FA is as low as noise level and no longer sensitive to cortical microstructural pattern. Other DTI-derived metrics such as MD have been used to characterize cortical microstructural changes in aging and Alzheimer’s disease (59, 60). A few more sophisticated dMRI-based models (e.g. 6–13) that usually require multi-shell dMRI can be used to study cortical internal microstructural complexity in the brain. The metrics derived from these models can be an effective and validated probe for quantifying cortical microstructure noninvasively. MK from DKI (7) that quantifies non-Gaussian diffusion occurring all over cerebral cortex was focused on in this study. As shown in Figures 3-7, the cortical MK is sensitive to inhomogeneous microstructural pattern across the cerebral cortex. On the other hand, as demonstrated in Figure 6b, DTI-derived FA measurements are low in nearly isotropic cortical regions and are not sensitive to microstructural changes across the cortical regions. Unlike FA that can be only used to delineate cortical microstructure in limited age range before or around birth, MK derived from DKI offers meaningful microstructural measurements in cerebral cortex of all age range in both human and animal brains. Figures 3-7 suggested that dMRI-based microstructural measurements have great potential to serve as an effective and noninvasive biomarker for assessing normal biological processes (e.g. brain development or aging) or detecting certain brain disorders.

### 4.2 Relationship between cortical MK and histological measurements

This study also revealed significant correlation between MK and ND across cortical regions with higher MK associated with higher ND. Two challenges were presented for MRI-histology correlations. The first challenge was to determine a reliable histological measurement. Given direct image intensity measurement based on digitized histological staining contrast can be susceptible to a variety of error sources, we adopted the structure tensor approach (30) to characterize fine fiber structures in the cerebral cortex and measure ND in the histological image. The pixel-wise structure tensor analysis used image partial derivatives as edge detectors and accurately detected anisotropic cortical fiber structures in the scale of sub-micrometer. Another challenge was a gap between histological image resolution and dMRI resolution for direct correlation between the dMRI-based measurements and histological measurements. To enable quantitative comparison between MK map and histology image, in this study an innovative approach was introduced. By blocking histological images into segments of 512×512 pixels equivalent to the scale of the upsampled DKI image resolution and defining ND in a novel way (right panel of Figure 1) at the same resolution as that of the upsampled MK map, histology-based ND measurements effectively validated the dMRI-based MK measurements. The significant linear correlation shown in Figure 7 revealed that the heterogeneity on dMRI-derived MK maps across cortical regions is consistent with the heterogeneity of ND from histology. This correlation suggested cortical MK could be a potential noninvasive surrogate biomarker for histology-based cortical ND in studies of brain development, aging and certain disorders where cortical microstructural changes are mainly associated with changes in ND.

### 4.3 Technical considerations, future directions and limitations

In this study, we optimized dMRI by changing resolution and b values for cortical MK measurements with sufficient SNR and anatomical details to delineate the microstructure within the thin cortical mantle of macaque brain. The dMRI sequences widely used for the brain scans in a 3T scanner have a typical resolution of 2 × 2 × 2 mm^3^. The dMRI sequence used in the present study has an in-plane resolution of 0.6×0.6mm^2^ (slice thickness 2mm) tailored specifically for relatively small macaque brains to reveal sufficient cortical microstructure details with a 3T scanner. With sufficient spatial resolution, the high-resolution MK map demonstrates clear heterogeneity of non-Gaussian diffusion properties across different cortical regions, ensuring accurate MK measurement and histology validation. Selection of appropriate b-values is especially important for kurtosis estimates in dMRI studies. By testing diffusion signal drop across a range of b values for *ex vivo* macaque brain samples, we successfully achieved the delicate balance between increasing the signal sensitivity to kurtosis and avoiding systemic errors in DKI metrics caused by overly high b-values (35). As demonstrated in Supporting Information Figure S1, two b-values (b=1500, 4500 s mm^−2^) were carefully selected. It is noteworthy that higher baseline b-value, namely 1500s mm^−2^ instead of widely used 1000s mm^−2^ in conventional in vivo DTI, was adopted in this study due to different diffusion properties of ex vivo brain tissue. Other b_max_ values adopted in kurtosis studies involving in *ex vivo* rat brains (61, 62) are lower than 4500s mm^−2^ used in this study. The differences of these b_max_ values are likely due to dramatic difference of cerebral cortical microstructural architecture (63, 64) between rats and macaques as well as factors such as different diffusion time Δ. Ex vivo scans with macaque brain samples immersed inside the fomblin significantly reduced the distortions caused by eddy currents and B0-inhomogeneity susceptibility (e.g. 65) despite relatively large b values applied in this study. For in vivo scans, more sophisticated alignment algorithms (e.g. 38) may need to be incorporated to improve alignment of DWIs with high b values.

As an exploratory study investigating the neuroanatomical underpinning of cortical MK measurements, there are several limitations including different macaque brain samples used for imaging from those used for histology, ROI matching between MK and ND map, relative small sample size, ex vivo instead of in vivo dMRI scans and lacking incorporation of soma contribution to MK. Overall consistent cortical microstructural profile among different macaque brains (e.g. Figure 4) suggests reproducibility of the MK heterogeneity pattern across different macaque brains and may justify correlation between MRI and histology with different samples. We also managed to match the anatomical ROIs in histology and those in MRI by rotating MRI and selecting MRI images demonstrating same anatomical landmarks shown in corresponding histological image. Despite the efforts for high-fidelity correspondence between the histological and MRI ROIs, subtle offsets of the ROI locations still existed and affected MK-ND correlation. Previous study (66) demonstrated that unlike absolute dMRI measurement such as mean diffusivity, relative dMRI measurements such as FA measurements were almost not affected in well-fixed postmortem brain tissues. In another study (67), MK of fixed brain WM regions increased significantly, but cortical MK increase in fixed cortex was much smaller. Cortical MK measurement from ex vivo MRI scans could still reflect cortical microstructure architecture in live brains. Due to scarcity of macaque brain samples, relatively small sample size of six ex vivo macaque brains was included in this study. Reproducibility test with larger sample size in future studies is warranted. To delineate neuroanatomical underpinning of cortical MK, we accounted for only fiber structures like axons or dendrites. Although fiber-like structure is a major contributor to MK measurement in the cerebral cortex (68-70), spherical structures such as somas may also contribute to MK. Future dMRI studies including advanced signal models and histology validation experiments incorporating signal contributions from both neurofilaments and somas are needed to delineate the contribution of each component to non-Gaussian diffusion in the cerebral cortex (68). In these future studies, different histological staining (e.g. Nissl staining) capable of delineating soma density will be added for validation purposes, besides the neurofilament staining. With the promising lead that cortical MK is sensitive to cortical ND demonstrated in this study, the ultimate goal is to develop a translational framework of “noninvasive neuropathology” that can quantify cortical neuronal and neurofilament density noninvasively based on dMRI. It is noteworthy that MK may potentially serve as a biomarker only when brain microstructural changes are mainly associated with neurofilament alterations. Specifically, for brain changes in development, aging and neurodegenerative diseases (e.g. Alzheimer’s disease and mild cognitive impairment) where microstructural alterations are largely associated with the neurofilaments, cortical MK may be sensitive to cortical ND changes associated with these brain processes. MK may not be correlated to ND in many other pathologies involving other significant microstructural alterations. For example, in acute ischemic stroke, MK increases significantly with the pathological events even though ND may decrease (e.g. 26, 71). The increased MK in ischemic lesion after acute stroke may result from the recruitment of inflammatory cells into the lesion regions (72). In a brain tumor study (73), increased MK was contributed by the abnormal proliferation of tumor cells observed with the increase grade of tumors.

## 5. Conclusion

In this study, we have delineated the cortical heterogeneity of MK with high resolution DKI of postmortem macaque brains and revealed significant linear relationship between cortical MK based on DKI and ND based on structure tensor analysis of histology image. This study revealed significant correlation between dMRI-based MK and histology-based ND within the cerebral cortical mantle, enabling assessment of histological measures in the cerebral cortex with noninvasive imaging techniques. The association of DKI metrics and histological features revealed in this study suggests great potential of dMRI-based microstructural measurement for identification of early microstructural biomarkers in the cerebral cortex in developmental, aging and diseased brains.

## Acknowledgments

This study was supported by NIH R01MH092535, NIH U54HD086984 and NIH R21EB009545.

## Data Availability Statement

The code and data that support findings of this study are openly available on GitHub. In-house diffusion kurtosis fitting code used in this study was made publicly available at https://github.com/ritaz0904/DK_fitting. Raw dMRI dataset from a representative macaque brain sample acquired in this study was made publicly available at https://github.com/ritaz0904/Macaque_DKI.

## Supporting Information

Additional Supporting Information may be found online in the Supporting Information section.

