## Supporting Information Figure S1 for "Neuroanatomical underpinning of diffusion kurtosis measurements in the cerebral cortex of healthy macaque brains"

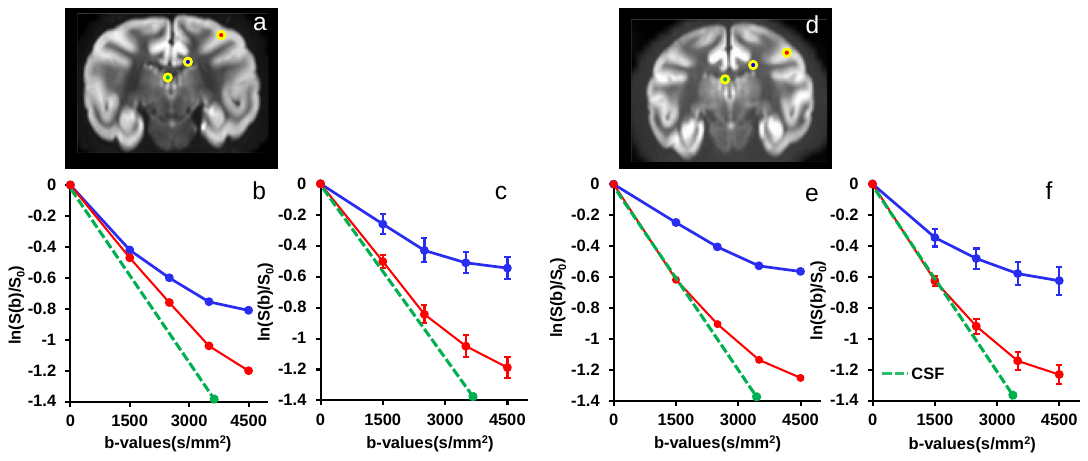

**Supporting Information Figure S1:** Reproducible diffusion signal drop at different b values in the postmortem macaque brain gray matter (GM), white matter (WM) and cerebrospinal fluid (CSF) for optimal b value selection for kurtosis estimation of the cerebral cortex. Panels a and d show GM, WM and CSF ROIs (red in GM, blue in WM and green in CSF) of two postmortem macaque brain samples. Panels b and e show consistent pattern of diffusion signal drop of GM (red line) and WM (blue line) at several b-values of 0, 1500, 2500, 3500 and 4500 s mm^-2^ with one diffusion gradent orientation. Signal drop of CSF (dashed green line) is shown as the reference. Panels c and f show consistesnt pattern of diffusion signal drop of GM (red line) and WM (blue line) averaged across 30 diffusion gradient orientations. At b=4500 s mm^-2^ clear non-Gaussian diffusion in cerebral cortex was captured, indicated by the deviation of the log (S(b)/S_0_) curve away from linear drop characterized by Gaussian diffusion typically observed in the CSF. S(b) and S_0_ are the signal with a certain b value and b of 0, respectively. Error bars indicate the standard deviation.

**Supporting Information Video S1:** Aligned diffusion-weighted images for macaque brain sample #1 across b-values 0, 1500s/mm2 and 4500s/mm2 after preprocessing. For each non-zero b value of 1500s/mm2 or 4500s/mm2, there were 30 diffusion gradient orientations.

**Supporting Information Video S2:** Aligned diffusion-weighted images for macaque brain sample #2 across b-values 0, 1500s/mm2 and 4500s/mm2 after preprocessing. For each non-zero b value of 1500s/mm2 or 4500s/mm2, there were 30 diffusion gradient orientations.

**Supporting Information Video S3:** Aligned diffusion-weighted images for macaque brain sample #3 across b-values 0, 1500s/mm2 and 4500s/mm2 after preprocessing. For each non-zero b value of 1500s/mm2 or 4500s/mm2, there were 30 diffusion gradient orientations.

**Supporting Information Video S4:** Aligned diffusion-weighted images for macaque brain sample #4 across b-values 0, 1500s/mm2 and 4500s/mm2 after preprocessing. For each non-zero b value of 1500s/mm2 or 4500s/mm2, there were 30 diffusion gradient orientations.

**Supporting Information Video S5:** Aligned diffusion-weighted images for macaque brain sample #5 across b-values 0, 1500s/mm2 and 4500s/mm2 after preprocessing. For each non-zero b value of 1500s/mm2 or 4500s/mm2, there were 30 diffusion gradient orientations.

**Supporting Information Video S6:** Aligned diffusion-weighted images for macaque brain sample #6 across b-values 0, 1500s/mm2 and 4500s/mm2 after preprocessing. For each non-zero b value of 1500s/mm2 or 4500s/mm2, there were 30 diffusion gradient orientations.
